# Processing bodies promote lysosomal quality control and cell survival during recovery from lysosomal damage

**DOI:** 10.1101/2025.08.27.672666

**Authors:** Jingyue Jia, Jacob Duran, Li Chen, Jing Pu, Pavel Ivanov

**Affiliations:** Center for Global Health, Department of Internal Medicine, University of New Mexico Health Sciences Center, Albuquerque, NM 87106, USA; Autophagy, Inflammation and Metabolism Center of Biochemical Research Excellence, Albuquerque, NM 87106, USA; Department of Molecular Genetics and Microbiology, University of New Mexico Health Sciences Center, Albuquerque, NM 87106, USA; Department of Medicine, Brigham and Women’s Hospital and Harvard Medical School; HMS Initiative for RNA Medicine, Boston, MA 02115, USA

## Abstract

Lysosomes are essential for cell survival but are highly susceptible to diverse physical and pathological stressors. Thus, the ability to initiate an acute damage response and promote recovery after stressor resolution is critical for maintaining cellular homeostasis and viability. Although recent studies have advanced our understanding of acute responses to lysosomal injury, the molecular mechanisms governing the recovery stage and distinguishing it from the acute phase remain poorly defined. Here, we delineate a key difference between these two stages in translational regulation and uncover lysosomal recovery from acute damage as a novel trigger for processing body (PB) formation. PBs are membraneless biomolecular condensates involved in RNA metabolism and translational reprogramming. We provide the first evidence that PBs are critical for lysosomal quality control and cell survival during recovery. Mechanistically, PBs are induced selectively during the recovery phase, but not during the acute damage response, through interactions with stress granules (SGs), distinct membraneless biomolecular condensates formed upon acute injury to stabilize damaged lysosomal membranes for repair. Functional analyses reveal that PBs promote lysosomal quality control by collaborating with SG-mediated membrane stabilization, while independently recruiting released cathepsins, thereby collectively supporting cell survival. Together, these findings establish PBs as central effectors of the lysosomal recovery program and underscore the broader relevance of biomolecular condensates in cellular responses to lysosomal damage and related disease processes.

## INTRODUCTION

Lysosomes are essential membrane-bound organelles that maintain cellular homeostasis. They function as digestive centers, breaking down a broad spectrum of intracellular and extracellular materials, and also act as signaling hubs that regulate metabolism, gene expression, quality control, and stress responses(Ballabio et al., 2020; Henn et al., 2025; Settembre et al., 2024). These functions expose lysosomes to various stressors, including biological, chemical, microbiological, environmental and mechanical agents, all of which can compromise membrane integrity(Jia et al., 2025; Meyer et al., 2024; Radulovic et al., 2025). Failure to promptly respond to such damage poses a lethal threat to cell survival, as the release of lysosomal contents such as cathepsins can trigger cell death and contribute to pathological consequences in health and disease (Jia et al., 2025; Lakpa et al., 2021; Radulovic et al., 2025).

Once lysosomes are exposed to stress and sustain damage, cells can elicit a sophisticated set of acute responses to cope with such stressful conditions(Jia et al., 2025). Mildly damaged lysosomes can be repaired through various mechanisms(Radulovic et al., 2022; Skowyra et al., 2018; Tan et al., 2022), while severely damaged lysosomes can be degraded by autophagy(Maejima et al., 2013). To compensate, lysosomal biogenesis and regeneration are upregulated(Bhattacharya et al., 2023; Napolitano et al., 2016). Meanwhile, cellular systems are remodeled to conserve resources and limit further damage, including the reprogramming of metabolic pathways(Jia et al., 2018; Jia et al., 2020a), alterations in endomembrane trafficking(Paddar et al., 2025; Wang et al., 2023), and a global shutdown of translation(Jia et al., 2022b). This translational shutdown can drive cellular phase separation, leading to the formation of stress granules (SGs)—reversible membraneless biomolecular condensates composed of translationally repressed RNAs and RNA-binding proteins(Jia et al., 2022b). SGs can stabilize damaged membranes(Bussi et al., 2023) and regulate translation to support cell survival during lysosomal stress(Duran et al., 2024; Ivanov et al., 2019). However, once the stress is resolved or the causative factors are removed, the mechanisms that restore damaged lysosomes to functional organelles, as well as the distinctions between these two stages, remain unclear.

Processing bodies (PBs) are another class of reversible membraneless biomolecular condensates formed via phase separation, driven by locally elevated densities of RNAs and proteins, similar to SGs(Riggs et al., 2020). PBs are also enriched in translationally repressed mRNAs but are uniquely defined by the presence of RNA-binding proteins involved in mRNA decapping, deadenylation, degradation, and translational repression(Luo et al., 2018; Standart et al., 2018). These include key decapping activators such as the RNA helicase DDX6 (DEAD-Box Helicase 6) and DCP1A (mRNA-Decapping Enzyme 1A)(Aizer et al., 2008; Minshall et al., 2009; Rzeczkowski et al., 2011; Serman et al., 2007). PB formation is considered important for mRNA metabolism, supporting mRNA storage(Brengues et al., 2005; Standart et al., 2018), degradation (Blake et al., 2024; Parker et al., 2007) as well as translational repression(Coller et al., 2005; Hubstenberger et al., 2017), thus contributing to translational reprogramming(Di Stefano et al., 2019; Ivanov et al., 2019).

Although both PBs and SGs are biomolecular condensates composed of RNAs and proteins, their formation dynamics and molecular composition reflect distinct roles in cellular stress responses(Riggs et al., 2020). PBs can exist under basal conditions and are further promoted by certain stressors, whereas SGs are exclusively stress-induced and uniquely contain specific translation initiation factors(Kedersha et al., 2005). Both granules sequester mRNAs, highlighting their shared involvement in RNA metabolism(Ivanov et al., 2019). However, SGs are thought to function primarily by transiently storing repressed mRNAs and stalled pre-initiation complexes, thereby effectively halting protein synthesis and promoting translational reprogramming(Riggs et al., 2020; Smith et al., 2024). In addition, SGs serve as signaling hubs by recruiting diverse signaling molecules to modulate stress responses(Kedersha et al., 2013; Thedieck et al., 2013; Wippich et al., 2013). In contrast, PBs are uniquely enriched in factors involved in mRNA decapping, deadenylation, degradation, and translational repression, implicating them in mRNA turnover and sustained translational silencing(Parker et al., 2007; Ripin et al., 2023). Despite these differences, PBs and SGs are compositionally and functionally interconnected, sharing both proteins and mRNA components(Kedersha et al., 2007). Shared factors such as DDX6 and UBAP2L (Ubiquitin Associated Protein 2 Like) contribute to the assembly of and interactions between SGs and PBs(Riggs et al., 2024; Ripin et al., 2024). Importantly, dysregulation of these granules has been implicated in various pathological contexts, including cancer progression, viral pathogenesis, and neurodegenerative disorders(Ripin et al., 2023). Together, these distinctions and shared features underscore the complementary and non-redundant roles of PBs and SGs in translational control and cellular adaptation to stress.

Previously, we found that acute lysosomal damage induces SG formation as a consequence of global translation shutdown(Jia et al., 2022b). This response is mediated by calcium release from damaged lysosomes, which promotes cell survival upon lysosomal injury(Duran et al., 2024). SGs have also been shown to play a direct role in physically plugging and stabilizing damaged lysosomal membranes to facilitate repair(Bussi et al., 2023). Although SGs have emerged as key players in the response to lysosomal damage, the role of PBs in this context remains unclear, despite their close association with SGs. In our earlier work, we did not observe increased PB formation during acute lysosomal damage(Jia et al., 2022b), suggesting that PBs are not part of the immediate stress response to lysosomal injury. However, given their close association with SGs and essential roles in cellular stress responses(Riggs et al., 2020), we hypothesized that PBs function primarily during the recovery phase, when translation is restored, to selectively facilitate stress resolution.

In this study, we found that translation is restored during the recovery stage of lysosomal damage—after the removal of lysosome-damaging agents—in contrast to the translational shutdown observed during the acute injury phase. We further discovered that PB formation is specifically promoted during the recovery phase, rather than during acute injury. This response was consistently observed across multiple cell types, including human macrophages. Notably, recovery-associated PB formation required both the presence of SGs and their interaction with SGs induced by acute damage. To assess the functional role of PBs in lysosomal recovery, we monitored lysosomal acidification and integrity in cells lacking a key PB component. These experiments revealed that impaired PB formation compromised the efficient restoration of lysosomal quality after acute injury, accompanied by increased cell death. Further studies showed that PBs associate with damaged lysosomes and SGs, suggesting that PBs and SGs may collaborate to stabilize injured lysosomes for repair. In addition, co-immunoprecipitation assays demonstrated increased interactions between PBs and cathepsins—but not SGs and cathepsins—during recovery, supporting an independent role of PBs in recruiting released cathepsins from damaged lysosomes to prevent cell death. Together, these findings reveal distinct patterns of protein translation between the acute and recovery stages of lysosomal damage, identify PBs as critical effectors of the cellular recovery program, and highlight their protective role in pathological contexts of lysosomal injury, where they may help mitigate damage-associated dysfunction.

## RESULTS

### Processing body formation is induced during recovery from lysosomal damage

Our previous study showed that PB formation remains unchanged during acute lysosomal damage(Jia et al., 2022b). In that experiment, we used DCP1A as a PB marker(Rzeczkowski et al., 2011) and treated U2OS cells (a human osteosarcoma epithelial cell line) with 2 mM Leu-Leu-O-Me (LLOMe), a well-characterized lysosome-damaging agent(Thiele et al., 1990), for 30 min. When we extended the LLOMe treatment to 1 h, it did not alter PB formation as assessed by confocal fluorescence microscopy (Fig. 1A). However, following 30 min of LLOMe treatment and a subsequent 30 min washout recovery period, we observed a marked increase in PB formation (Fig. 1A). To quantify this increase, we employed high-content microscopy (HCM) using two additional PB-specific markers, LSM14A and DDX6(Ayache et al., 2015) and observed a significant increase in PBs (Figs. 1B and 1C). HCM is a machine-assisted, high-throughput imaging approach that enables analysis of large cell populations, providing quantitative data and statistical outputs(Claude-Taupin et al., 2021; Gu et al., 2019; Jia et al., 2018; Jia et al., 2020a; Jia et al., 2020b; Jia et al., 2022b; Wang et al., 2023). In parallel, we monitored lysosomal acidification using a LysoTracker staining assay(Jia et al., 2020b; Jia et al., 2022b), which showed a significant reduction in acidification during LLOMe treatment, followed by recovery during the washout phase (Fig. 1D). Notably, treatment with a higher dose of LLOMe, which induces more severe lysosomal damage, led to more robust PB formation during the recovery phase (Fig. S1A). In contrast, extending the recovery period after LLOMe washout resulted in a gradual decline in PB formation (Fig. S1B), accompanied by increased LysoTracker puncta staining, indicating progressive recovery of lysosomal acidification (Fig. S1C). These findings suggest that PB formation is linked to the extent of lysosomal injury, pointing to a potential role for PBs in facilitating lysosomal recovery from damage. Moreover, we observed robust PB formation during the recovery phase following LLOMe-induced lysosomal damage in additional cell types, including Huh7 cells (a human hepatocellular carcinoma cell line) and human primary macrophages (Figs. S1D and S1E). While lysosomal damage is known to trigger SG formation as a key acute response to lysosomal injury(Bussi et al., 2023; Duran et al., 2024; Jia et al., 2022b), our findings indicate that PB formation is a distinct recovery-associated event (Fig. 1E). Together, these results support a model in which SGs respond to acute lysosomal stress, whereas PBs emerge during the resolution phase, potentially facilitating cellular recovery from lysosomal damage (Fig. 1E).

**Figure 1.**
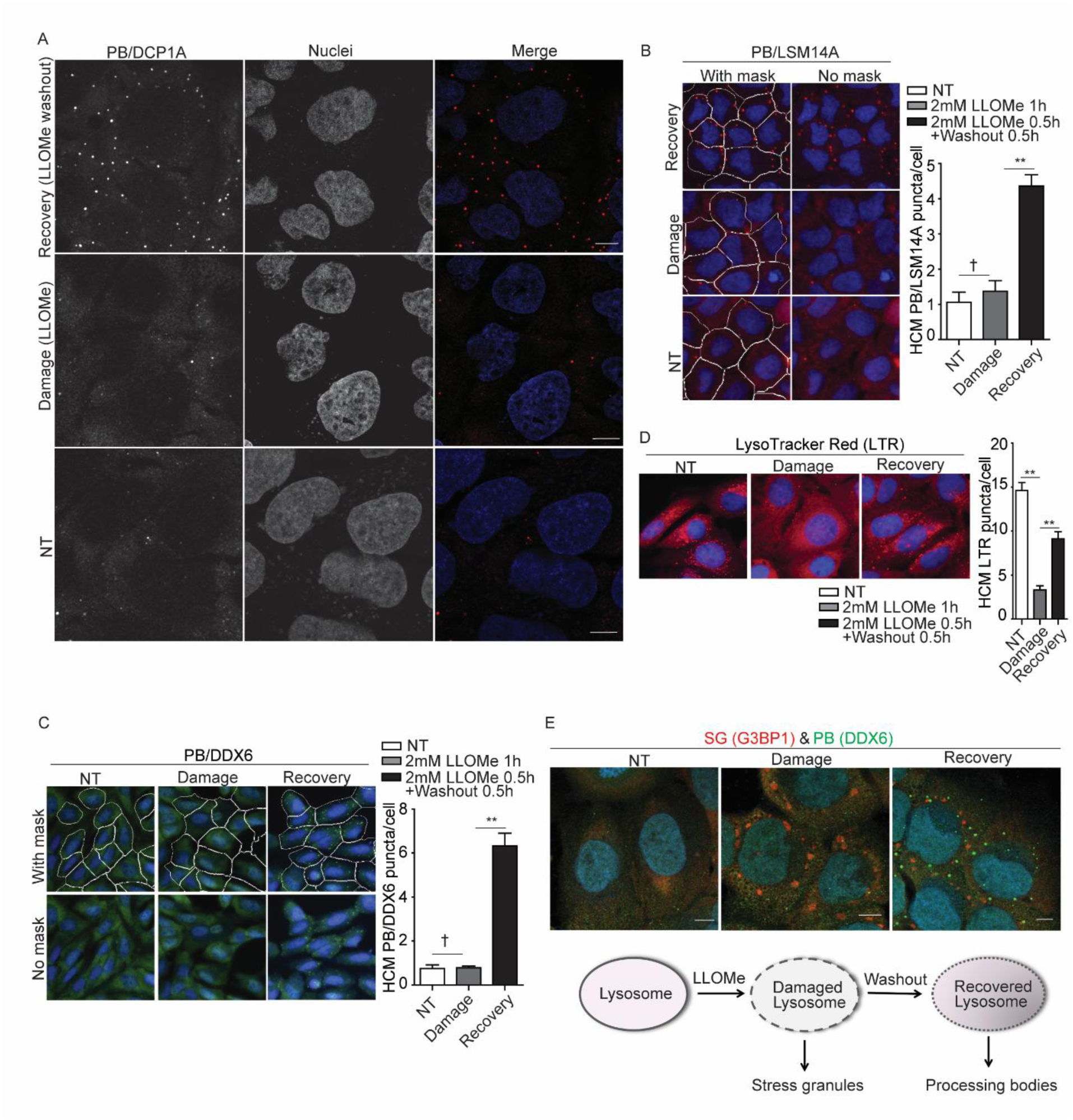
Processing body formation is induced during recovery from lysosomal damage. **(A)** Confocal microscopy analysis of the processing body (PB) marker DCP1A (Alexa Fluor 568) in U2OS cells treated with 2 mM LLOMe for 1 h (damage condition), or for 30 min followed by a 30 min recovery after washout (recovery condition). Scale bar, 10 μm. **(B)** Quantification of the PB marker LSM14A by high-content microscopy (HCM) in U2OS cells treated with 2 mM LLOMe for 1 h to induce damage, or for 30 min followed by a 30 min recovery after washout. White masks, algorithm-defined cell boundaries; red masks, computer-identified LSM14A puncta. **(C)** Quantification of the PB marker DDX6 by HCM in U2OS cells treated with 2 mM LLOMe for 1 h to induce damage, or for 30 min followed by a 30 min recovery after washout. White masks, algorithm-defined cell boundaries; green masks, computer-identified DDX6 puncta. **(D)** Status of acidified organelles in U2OS cells assessed by LysoTracker (LTR) staining and HCM during lysosomal damage (2 mM LLOMe for 1 h) or recovery (2 mM LLOMe for 30 min followed by a 30 min recovery after washout). Red masks, computer-identified LTR puncta. **(E)** Confocal microscopy analysis of the stress granule (SG) marker G3BP1 (Alexa Fluor 568) and the PB marker DDX6 (Alexa Fluor 488) in U2OS cells treated with 4 mM LLOMe for 1 h (damage condition), or for 30 min followed by a 30 min recovery after washout (recovery condition). Scale bar, 10 μm. Schematic summary of the findings in Figure 1 and S1. NT, untreated cells. Data, means ± SEM (n = 3); HCM: n ≥ 3 (each experiment: 500 valid cells per well, ≥5 wells/sample). † p ≥ 0.05 (not significant), **p < 0.01, ANOVA. See also Figure S1.

### Processing body formation during lysosomal damage recovery requires stress granules

We next investigated the potential mechanisms regulating PB formation during lysosomal damage recovery. Since SGs are initially induced by acute lysosomal stress, and mounting evidence indicates that SGs and PBs interact through shared components(Buchan, 2024; Ivanov et al., 2019; Riggs et al., 2024; Riggs et al., 2020; Ripin et al., 2024), we hypothesized that PB formation may be regulated by SGs. SG formation is driven by global translation shutdown mediated by phosphorylation of eIF2α (eukaryotic translation initiation factor 2 Subunit Alpha)(Duran et al., 2024; Kedersha et al., 1999). We and others have previously observed eIF2α phosphorylation and translation shutdown during acute lysosomal damage(Bussi et al., 2023; Duran et al., 2024; Jia et al., 2022b). We therefore examined whether recovery from lysosomal damage reverses eIF2α phosphorylation and restores global protein translation. Indeed, phosphorylation returned to baseline levels after a 30 min recovery period (Fig. 2A). Using a puromycin incorporation assay, we found that global translation gradually resumed following LLOMe washout, requiring over 1 h of recovery to return to steady-state levels (Fig. 2B). Interestingly, protein synthesis exceeded baseline levels after 1.5 h of recovery (Fig. 2B), which may reflect an elevated demand for protein production as cells work to restore homeostasis. In parallel, we assessed the protein levels of key components of PBs and SGs during the recovery phase. DDX6, a core factor essential for PB formation(Minshall et al., 2009; Serman et al., 2007), remained stable throughout the time course (Fig. 2A). In contrast, the protein levels of G3BP1 and G3BP2 —critical for SG assembly(Yang et al., 2020)—gradually decreased over time (Fig. 2A), suggesting SG disassembly or clearance. To test this, we quantified SG puncta using the SG-specific marker EIF4G(Ivanov et al., 2019) and HCM. As expected, SG puncta progressively decreased with longer recovery durations (Fig. 2C).

**Figure 2.**
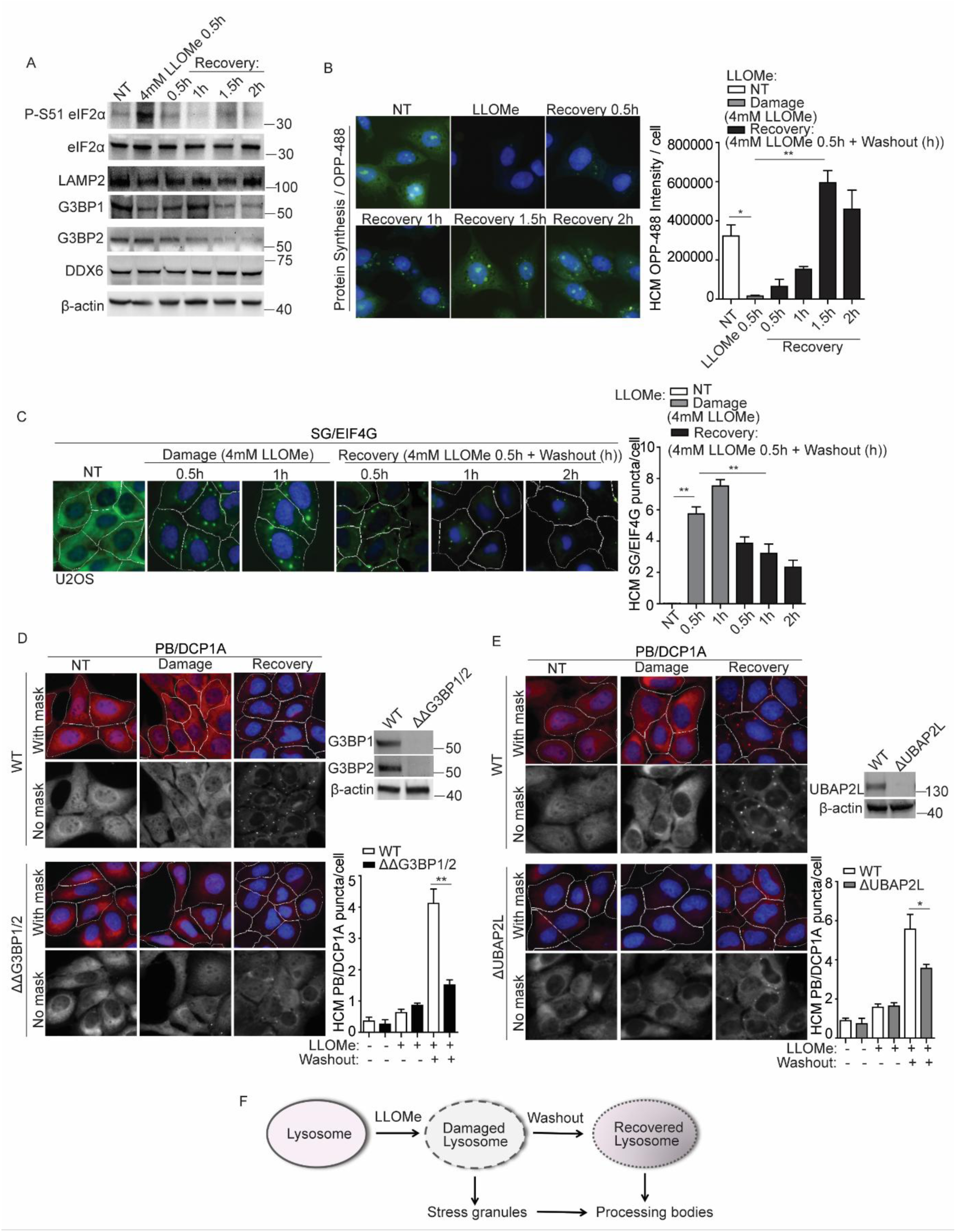
Processing body formation during lysosomal damage recovery requires stress granules. **(A)** Immunoblot analysis of eIF2α phosphorylation at S51 and other indicated proteins in U2OS cells treated with 4 mM LLOMe for 30 min to induce damage, or treated for 30 min followed by a time-dependent recovery after washout. **(B)** HCM analysis of protein synthesis in U2OS cells using the Click-iT Plus OPP Alexa Fluor 488 Protein Synthesis Assay (Thermo Fisher Scientific), following treatments described in (A). **(C)** Quantification of the stress granule (SG) marker EIF4G by HCM in U2OS cells treated with 4 mM LLOMe for 30 min or 1 h to induce damage, or with 4 mM LLOMe for 30 min, followed by a time-dependent recovery. White masks, algorithm-defined cell boundaries; green masks, computer-identified EIF4G puncta. **(D)** Quantification of the PB marker DCP1A by HCM in U2OS wildtype (WT) and G3BP1&2 double knockout (ΔΔG3BP1/2) cells treated with 2 mM LLOMe for 1 h to induce damage, or for 30 min followed by a 30 min recovery after washout. White masks, algorithm-defined cell boundaries; red masks, computer-identified DCP1A puncta. No mask image is shown in grayscale. **(E)** Quantification of the PB marker DCP1A by HCM in U2OS wildtype (WT) and UBAP2L knockout (ΔUBAP2L) cells treated with 2 mM LLOMe for 1 h to induce damage, or for 30 min followed by a 30 min recovery after washout. White masks, algorithm-defined cell boundaries; red masks, computer-identified DCP1A puncta. No mask image is shown in grayscale. **(F)** Schematic summary of the findings in Figure 2 and S2. NT, untreated cells. Data, means ± SEM (n = 3); HCM: n ≥ 3 (each experiment: 500 valid cells per well, ≥5 wells/sample). *p < 0.05, **p < 0.01, ANOVA. See also Figure S2.

Since SGs dissipate and PBs emerge during lysosomal recovery, we next tested whether SGs are required for PB formation in this context. First, we used SG-deficient U2OS cells with genetic deletions of both G3BP1 and G3BP2 (ΔΔG3BP1/2)(Kedersha et al., 2016). In these cells, PB formation during the recovery stage was significantly inhibited compared to wildtype cells (Fig. 2D). As expected, SG formation during LLOMe treatment was abolished in ΔΔG3BP1/2 cells (Fig. S2A). This suggests that SGs are necessary for subsequent PB formation during recovery. To further validate this, we knocked down G3BP1 and G3BP2 in Huh7 cells, which similarly reduced PB formation during recovery and abolished SG formation during LLOMe treatment (Figs. S2B-G). We also applied G3BP inhibitors FAZ3532 or FAZ3780, which bind to the dimerization domain of G3BP1/2, and specifically disrupt G3BP-RNA co-condensation and the SG network(Freibaum et al., 2024). Consistently, these inhibitors prevented PB formation during the recovery phase while inhibiting SG formation during LLOMe treatment (Figs. S2B-G). Furthermore, we employed UBAP2L-deficient cells (ΔUBAP2L), which lack UBAP2L—a scaffolding factor known to mediate the functional association between PBs and SGs(Riggs et al., 2024). In these cells, PB formation during recovery was impaired (Fig. 2E), and SG formation during the acute response phase was also significantly reduced (Fig. S2A). These findings indicate that SG formation and their communication with PBs are essential for PB assembly during lysosomal damage recovery.

To further strengthen this result, we used BAPTA-AM, a calcium chelator, to block calcium signaling from damaged lysosomal required for SG formation, as we reported previously(Duran et al., 2024). This treatment abolished SGs during acute lysosomal damage and consequently prevented PB formation during the recovery stage (Figs. S2H and I), highlighting the dependence of PB assembly on SGs. Moreover, treatment with the V-ATPase inhibitor Bafilomycin A1(Hooper et al., 2021), which disrupts lysosomal acidification, did not induce the formation of SGs and PBs at either stage (Figs. S2H and I), indicating that lysosomal membrane damage—but not lysosomal neutralization—is required for SG and PB formation. Altogether, these data support a signaling pathway in which PB formation during lysosomal recovery depends on SGs induced by acute lysosomal damage (Fig. 2F).

### Processing bodies and stress granules promote lysosomal quality recovery from damage

Since SGs have been shown to play an important role in lysosomal quality control by stabilizing damaged lysosomal membranes for repair(Bussi et al., 2023), we asked whether PB formation is also critical for maintaining lysosomal quality during recovery. To address this question, we employed U2OS DDX6 knockout (DDX6KO) cells, which lack DDX6—an essential factor required for PB formation(Ayache et al., 2015; Ripin et al., 2024; Serman et al., 2007). We first confirmed that DDX6 is critical for PB formation during lysosomal recovery, as DDX6KO cells failed to form PBs during the LLOMe washout phase (Fig. 3A). Despite this defect, DDX6KO cells showed no change in the expression levels of the lysosomal membrane protein LAMP2(Eskelinen, 2006) or lysosome-enriched proteases such as cathepsins(Repnik et al., 2012) compared to wildtype cells under resting conditions (Fig. S3A). This was further reflected by comparable steady-state lysosomal acidification between wildtype and DDX6KO cells, as measured by LysoTracker staining (Fig. 3B). We then evaluated lysosomal acidification during the recovery stage. As previously observed, wildtype cells restored lysosomal acidification following LLOMe washout (Fig. 3B). However, DDX6KO cells failed to re-establish acidification during this phase (Fig. 3B). To further investigate this effect, we examined lysosomal acidification in G3BPs or UBAP2L knockout cells, which lack SG formation or SG–PB interactions—both upstream events in PB formation during lysosomal recovery. We first assessed the contributions of DDX6, G3BPs, and UBAP2L to PB or SG formation. DDX6 played a dominant role in PB formation (Fig. S3B), while G3BPs were essential for SG formation (Fig. S3C), consistent with their roles under other stress conditions(Minshall et al., 2009; Yang et al., 2020). Although PBs were not completely abolished in G3BPs and UBAP2L knockout cells, their formation following lysosomal damage recovery was significantly reduced, underscoring their roles as key upstream regulators of PB formation. Importantly, the loss of these key PB regulators also impaired lysosomal recovery, as evidenced by reduced LysoTracker staining in both G3BPs and UBAP2L knockout cell lines (Fig. S3D). This defect was further validated using Glycyl-L-phenylalanine 2-naphthylamide (GPN), another well-established lysosomal damaging reagent(Berg et al., 1994; Thurston et al., 2012). After GPN-induced damage, PBs were again induced in wildtype cells but remained absent in DDX6KO cells (Fig. S3E), and lysosomal acidification was restored only in wildtype cells (Fig. S3F). Collectively, these findings highlight the essential role of PB formation in lysosomal recovery from damage.

**Figure 3.**
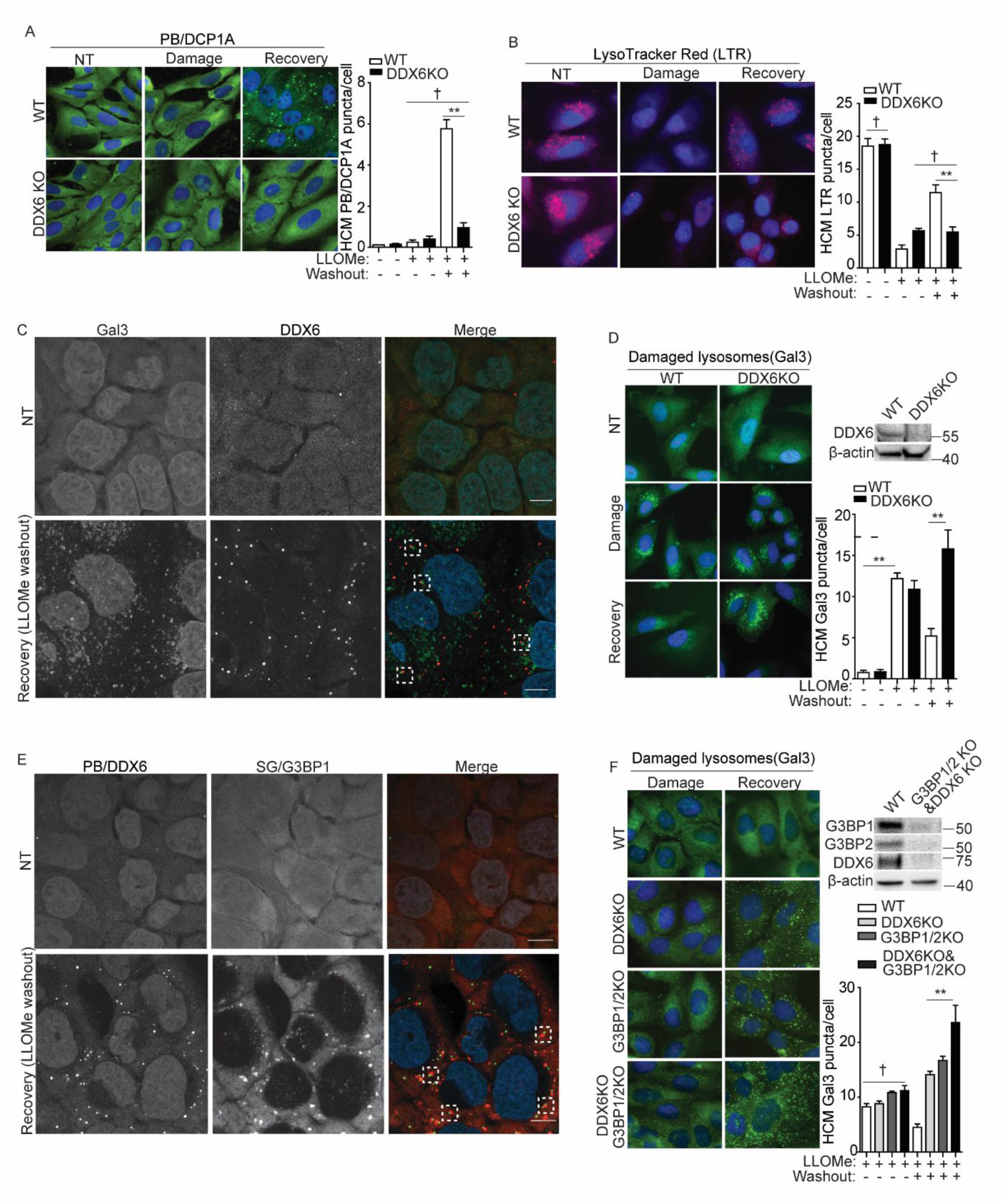
Processing bodies and stress granules promote lysosomal quality recovery from damage. **(A)** Quantification of the PB marker DCP1A by HCM in U2OS wildtype (WT) and DDX6 knockout (DDX6KO) cells treated with 2 mM LLOMe for 1 h to induce damage, or for 30 min followed by a 30 min recovery after washout. **(B)** Status of acidified organelles in U2OS wildtype (WT) and DDX6 knockout (DDX6KO) cells assessed by LysoTracker (LTR) staining and HCM during lysosomal damage (2 mM LLOMe for 1 h) or recovery (2 mM LLOMe for 30 min followed by a 30 min recovery after washout). **(C)** Confocal microscopy analysis of U2OS cells for the PB marker DDX6 (Alexa Fluor 568) and the damaged lysosome marker galectin-3 (Gal3) (Alexa Fluor 488) after treatment with 2 mM LLOMe for 1 h to induce damage, or for 30 min followed by a 30 min recovery after washout. Scale bar, 10 μm. **(D)** Quantification of the Gal3 by HCM in U2OS wildtype (WT) and DDX6 knockout (DDX6KO) cells treated with 2 mM LLOMe for 1 h to induce damage, or for 30 min followed by a 120 min recovery after washout. **(E)** Confocal microscopy analysis of U2OS cells for the PB marker DDX6 (Alexa Fluor 488) and the SG marker G3BP1 (Alexa Fluor 568) after treatment with 4 mM LLOMe for 1 h to induce damage, or for 30 min followed by a 30 min recovery after washout. Scale bar, 10 μm. **(F)** Quantification of the Gal3 by HCM in wildtype (WT), PB-deficient (DDX6KO), SG-deficient (G3BP1/2KO) or PB-SG double-deficient (G3BP1/2KO & DDX6KO) cells treated with 2 mM LLOMe for 30 min to induce damage, or for 30 min followed by a 120 min recovery after washout. White masks, algorithm-defined cell boundaries; green masks, computer-identified Gal3 puncta. Data, means ± SEM (n = 3); HCM: n ≥ 3 (each experiment: 500 valid cells per well, ≥5 wells/sample). † p ≥ 0.05 (not significant), **p < 0.01. See also Figure S3.

Since PBs can promote lysosomal recovery within a short 30 minute recovery period, this prompted us to investigate their direct role in damaged lysosomes. Given that SGs have been shown to stabilize damaged lysosomal membranes and promote repair(Bussi et al., 2023), and PBs share similar properties as biomolecular condensates with SGs(Banani et al., 2017), we asked whether PBs also facilitate lysosomal recovery through a comparable mechanism. Supporting this possibility, PBs were found to associate with damaged lysosomes labeled by galectin-3 (Gal3) (Fig. 3C), a well-established marker of luminal glycans exposure on compromised lysosomes(Aits et al., 2015). Consistently, PB deficiency markedly impaired lysosomal recovery compared to wildtype, resulting in an increased number of damaged lysosomes during the recovery phase (Fig. 3D), in line with reduced lysosomal acidification under these conditions (Fig. 3B). PBs and SGs have previously been reported to form physical associations under stress (Kedersha et al., 2005; Sanders et al., 2020; Souquere et al., 2009). To investigate whether such interactions occur during lysosomal recovery, we examined PB-SG colocalization using confocal fluorescence microscopy and HCM. PBs and SGs were frequently associated during the recovery phase (Fig. 3E), with approximately 30-40% spatial overlap (Fig. S3G). These findings suggest that PBs may support lysosomal recovery either in cooperation with or independently of SGs, with their independent role potentially compensating for the reduced presence of SGs during the recovery stage (Fig. 2C). Therefore, the presence of both PBs and SGs is important for lysosomal recovery. This was further supported by the impaired recovery capacity and increased accumulation of damaged lysosomes in PB-deficient (DDX6KO), SG-deficient (G3BP1/2KO), and PB-SG double-deficient (G3BP1/2KO & DDX6KO) cells (Ripin et al., 2024) compared with wildtype during the recovery stage (Fig. 3F). The deficiency of both PB and SG formation was confirmed in PB-SG double-deficient cells under conditions of acute damage as well as during recovery (Figs. S4A, B). Together, these results suggest that PB and SG formation are both critical, and at least partially cooperative, components of the lysosomal quality control system.

### Processing body formation supports cell survival during lysosomal damage recovery

Previously, we showed that SG formation is important for promoting cell survival during lysosomal damage(Duran et al., 2024; Jayabalan et al., 2025). Here, we investigated whether PB formation also contributes to cellular survival under lysosomal stress. We first measured cell death in wildtype and DDX6KO U2OS cells during acute lysosomal damage (1 h LLOMe treatment) and during the recovery phase (30 min LLOMe treatment followed by a 30 min washout recovery). Cell death was assessed using both a lactate dehydrogenase (LDH) release assay, which measures non-specific membrane leakage(Kimura et al., 2017), and live-cell viability via AMNIS imaging flow cytometry(Duran et al., 2024; Wang et al., 2023). Notably, cell death was significantly increased in DDX6KO cells during the recovery phase, but not during acute damage (Figs. 4A, B). To validate these results, we extended our analysis to Huh7 cells. Knockdown of DDX6 (DDX6^KD^) in Huh7 cells resulted in a marked reduction in PB formation during the GPN washout phase (Fig. S4C). Consistent with observations in U2OS cells, DDX6^KD^ Huh7 cells exhibited significantly increased cell death following GPN washout, but not during acute damage (Figs. S4D, E), underscoring the critical role of PBs in lysosomal recovery. Importantly, re-expression of DDX6 in DDX6KO cells restored PB formation upon LLOMe washout (Fig. 4C) and significantly reduced cell death (Fig. 4D), underscoring the importance of PBs in supporting cell survival.

**Figure 4.**
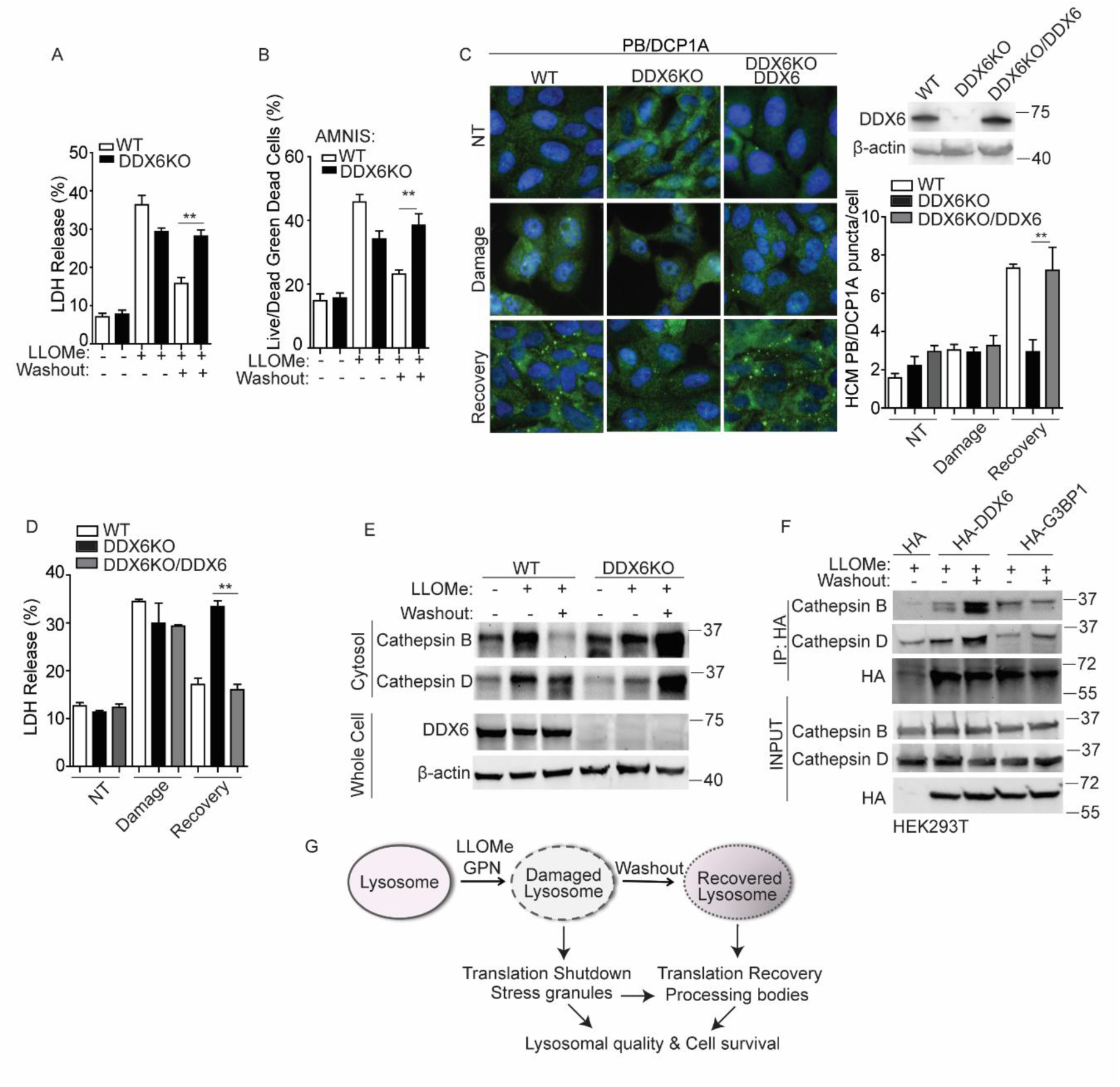
Processing body formation supports cell survival during lysosomal damage recovery. (**A**) Cell death analysis of supernatants from U2OS wildtype (WT) and DDX6 knockout (DDX6KO) cells by an LDH release assay. Cells were treated 2 mM LLOMe for 1 h to induce damage or treated for 30 min followed by a 30 min recovery after washout. (**B**) Quantification of cell death using the AMNIS system and Live/Dead^TM^ stain kit in U2OS wildtype (WT) and DDX6 knockout (DDX6KO) cells. Cells were treated as described in (A) and then stained using Live/Dead^TM^ stain kit (ThermoFisher). **(C)** Quantification of the PB marker DCP1A (Alexa Fluor 488) by HCM in U2OS wildtype (WT), DDX6 knockout (DDX6KO) with or without DDX6 overexpression. Cells were treated with 2 mM LLOMe for 1 h to induce damage or treated for 30 min followed by a 30 min recovery after washout. (**D**) Cell death analysis of supernatants from U2OS wildtype (WT) and DDX6 knockout (DDX6KO) cells with or without DDX6 overexpression by a LDH release assay. Cells were treated as described in (C). **(E)** Immunoblot analysis of cathepsins in the cytosol of U2OS wildtype (WT) and DDX6 knockout (DDX6KO) cells treated with 2 mM LLOMe for 1 h to induce damage or treated for 30 min followed by a 30 min recovery after washout. **(F)** Co-immunoprecipitation analysis of interactions between HA-DDX6/G3BP1 and cathepsins in HEK293T cells. Cells transfected with HA control or HA-DDX6/G3BP1 were treated with 1 mM LLOMe for 1 h or 30 min followed by a 30 min recovery. Cell lysates were immunoprecipitated with anti-HA antibody and immunoblotted for indicated proteins. **(G)** Schematic summary of the findings in this study. NT, untreated cells. Data, means ± SEM (n = 3); HCM: n ≥ 3 (each experiment: 500 valid cells per well, ≥5 wells/sample). **p < 0.01, ANOVA. See also Figure S4.

Given that the release of lysosomal contents such as cathepsins B and D can trigger cell death(Boya et al., 2003; Deiss et al., 1996; Eriksson et al., 2020; Johansson et al., 2003; Zhu et al., 2020), we next examined cytosolic cathepsin release(Appelqvist et al., 2011; Eriksson et al., 2020) in DDX6KO U2OS cells. As expected, LLOMe treatment induced cathepsin release in both wildtype and DDX6KO cells (Fig. 4E). However, during the washout phase, cytosolic cathepsins levels remained elevated in DDX6KO cells relative to wildtype (Fig. 4E)—reflecting impaired lysosomal recovery. The accumulation of cytosolic cathepsins in PB-deficient cells may result from defective lysosomal quality control or additional mechanisms. Given the reported role of SGs in recruiting proteins to rewire cellular signaling(Kedersha et al., 2013), we tested whether PBs may similarly recruit cytosolic cathepsins. Interestingly, immunopurification of PBs using HA-DDX6 and of SGs using HA-G3BP1 both revealed interactions with cathepsins B and D during acute damage (Fig. 4F). However, during the recovery stage, only PBs—not SGs—showed increased association with cathepsins (Fig. 4F). These findings suggest that PBs may recruit cathepsins released from damaged lysosomes, thereby suppressing cell death.

Acute lysosomal damage triggers SG formation, while recovery promotes PB assembly—two biomolecular condensates with interdependent dynamics that play key roles in regulating cell fate during stress(Bussi et al., 2023; Duran et al., 2024; Jayabalan et al., 2025; Riggs et al., 2020). To further elucidate their roles in cellular fate under lysosomal stress, we utilized PB-SG double-deficient cells (Ripin et al., 2024). Loss of both SGs and PBs significantly increased cell death during both acute damage and recovery (Fig. S4F), underscoring their importance in the cellular response to lysosomal damage. Collectively, our findings identify a novel mechanism by which how cells respond to lysosomal damage. Upon acute damage, translation is shut down and SGs form, whereas during the recovery phase, translation resumes, SGs dissipate and PB formation is triggered in an SG-dependent manner; PBs may collaborate with SGs or act independently to support cell survival by promoting lysosomal quality control or recruiting released cathepsins (Fig. 4G). Thus, SGs and PBs function as essential and cooperative components that maintain cellular homeostasis throughout the entire course of the response to lysosomal damage.

## DISCUSSION

n this study, we identify translation restoration during the recovery phase of lysosomal damage— distinct from the translational shutdown induced by acute injury—and establish PB formation as a key event in lysosomal recovery, providing new insights into the cellular homeostatic response to lysosomal stress. We further uncover the regulatory mechanisms driving PB formation during the recovery phase and clarify its functional roles in this process. We show that PB formation is specifically promoted during lysosomal recovery rather than during acute stress, and that this recovery-phase PB formation requires interactions with SGs initiated during the acute damage phase that stabilize damaged lysosomal membranes. Critically, PB formation is essential for restoring lysosomal integrity and ensuring cell survival following damage, either through its collaboration with SGs or by independently recruiting released lysosomal contents, thereby establishing PBs as integral components of the lysosomal recovery process. Together with previous studies demonstrating the role of SGs in the acute response to lysosomal damage(Bussi et al., 2023; Duran et al., 2024; Jayabalan et al., 2025; Jia et al., 2022b), this work highlights a fundamental interplay between lysosomes and biomolecular condensates in orchestrating cellular adaptation to stress.

PB formation predominantly occurs during the lysosomal recovery phase, whereas SG formation is more prominent during acute lysosomal damage. This difference is largely driven by the distinct regulatory factors that govern their assembly across the two stages. SG formation is triggered by cellular stress that stalls translation initiation, leading to a sudden accumulation of messenger ribonucleoprotein particles (mRNPs), which rapidly condense into discrete foci through networks of protein and RNA interactions(Ivanov et al., 2019; Kedersha et al., 1999). Thus, acute lysosomal stress induces a surge of translationally repressed mRNAs that drives SG assembly(Jia et al., 2022b). Upon lysosomal recovery, the global translation shutdown is relieved, leading to restored protein synthesis and the disassembly or clearance of SGs, as shown in Figure 2A–C. In contrast, PB formation occurs independently of global translation shutdown and is primarily regulated by alterations in mRNA metabolism, such as enhanced mRNA deadenylation(Ivanov et al., 2019; Kedersha et al., 2005; Luo et al., 2018). While PBs also contain translationally repressed RNAs, they are uniquely enriched in proteins involved in diverse post-transcriptional processes, including mRNA degradation, translational control, and RNA interference(Blake et al., 2024; Parker et al., 2007; Standart et al., 2018; Teixeira et al., 2005). Thus, recovery-induced PB formation may reflect changes in mRNA metabolism, such as increased mRNA deadenylation during the recovery stage compared with acute damage. In addition, the dependence of PB formation on SGs, together with the reduction of SGs during recovery, suggests a potential transfer of RNA or protein components from SGs to PBs, pointing to a novel form of PB–SG crosstalk. Altogether, differences in global transcription and translation between the acute damage and recovery stages, triggered by lysosomal status, may drive the distinct temporal appearance of SGs and PBs to maintain lysosomal homeostasis.

This temporal coordination reflects their distinct biological roles and represents a strategic cellular response to maximize survival under stress. SGs formed in response to acute damage can promote cell survival by reprogramming gene expression—separating silenced mRNAs from active translation sites to conserve energy and resources, activating stress response pathways(Ivanov et al., 2019; Jia et al., 2022a; McCormick et al., 2017), and physically plugging damaged lysosomes to facilitate membrane repair(Bussi et al., 2023). Since SGs are reduced during lysosomal recovery, they appear to function primarily in the acute phase of lysosomal damage. In addition, our results show that PBs influence lysosomal quality and cell survival within a relatively short time frame—detectable as early as 30 minutes after washout recovery— indicating a novel function distinct from their established roles in RNA metabolism, which may occur later. Importantly, we found that PBs colocalize with SGs and are required for lysosomal quality control, potentially compensating for the reduced presence of SGs in stabilizing damaged lysosomes. Moreover, the increased association of lysosomal contents such as cathepsins with PBs during recovery highlights their potential role in sequestering harmful factors to prevent cell death. Similar to SGs, these findings suggest that PBs can act as signaling hubs, modulating stress-response pathways and promoting lysosomal recovery by sequestering key regulatory components. Moreover, since PBs are known to serve as sites for mRNA degradation, storage, and translational repression (Blake et al., 2024; Brengues et al., 2005; Coller et al., 2005; Cougot et al., 2004; Sheth et al., 2003; Standart et al., 2018), they may also exert additional functions in lysosomal recovery over longer time courses beyond those identified in this study. PBs may facilitate the selective translation of high-priority mRNAs by degrading, repressing, or storing nonessential transcripts, possibly through the redistribution of stalled mRNAs released during SG disassembly. They may also sequester damaged RNAs generated during lysosomal stress, protecting the cell from their potential toxicity. Together, the formation and dissolution of PBs and SGs constitute a novel, coordinated, and dynamic system in response to lysosomal damage, opening new avenues for investigating the molecular mechanisms of lysosomal damage responses and the potential roles of PBs and SGs in these processes.

Although PBs and SGs are closely linked biomolecular condensates involved in translational control, their composition, interactions, and functions appear to be context-dependent(Riggs et al., 2020). For example, both PBs and SGs increase following arsenite exposure, whereas heat shock induces SGs without significantly affecting PBs(Kedersha et al., 2005). Similarly, both our previous work and the present study show that acute lysosomal damage triggers SG formation but does not induce PBs under our testing conditions(Jia et al., 2022b). Nevertheless, we cannot rule out the possibility that differences in lysosomal stressors or cell type may also influence PB formation during the acute response, since the ability of cells to cope with damage can vary. In general, lysosomal damage causes translational shutdown and SG assembly, and if cells are subsequently able to recover from such stress, with translation resumption and SG disassembly or clearance, PB formation is then promoted. Interesting, an increased trend of LysoTracker staining was observed under low-dose LLOMe treatment, suggesting that cells may partially restore lysosomal function even in the continued presence of the damaging reagent(Radulovic et al., 2018). This indicates that cells can overcome low-level stress to recover damaged lysosomes despite ongoing insult, and that the roles of translational regulation, SGs, and PBs in this context warrant further investigation. Thus, the distinct behavior of PBs and SGs under specific conditions may reflect different needs for translational control, highlighting the complexity of cellular transcriptomic and proteomic responses to environmental cues.

This study uncovers two critical stages of cellular responses to lysosomal damage—the acute damage phase and the recovery phase—and identifies their major differences, which are highly relevant to both physiological and pathological conditions associated with lysosomal dysfunction. Using pharmacological agents to mimic these stages, we reveal distinct translational states and dynamic SG–PB interactions. In a disease context, such features could serve as markers to distinguish between acute lysosomal damage and recovery, and may hold potential as therapeutic indicators. Furthermore, our findings pave the way for future investigations into how SG–PB pathway intersects with broader lysosomal damage responses and influences disease outcomes. For example, recent studies have shown that both PBs and p62 (Sequestosome-1) contribute to inflammasome regulation(Barrow et al., 2024), suggesting a potential link between PBs and broader lysosomal damage responses, given that both p62 and inflammasomes are closely associated with lysosomal damage(Hornung et al., 2008; Jia et al., 2025; Meyer et al., 2024; Yang et al., 2023). Notably, PBs, SGs and lysosomal stress are increasingly recognized as central players in viral infection, neurodegenerative diseases, and cancer(Ivanov et al., 2019; Jia et al., 2025; Lakpa et al., 2021; Riggs et al., 2020). Moreover, lysosomal stress has emerged—as demonstrated here and in prior studies—as a physiologically and pathologically relevant trigger for PB and SG formation(Bussi et al., 2023; Duran et al., 2024; Jayabalan et al., 2025; Jia et al., 2022b). A deeper understanding of their interplay will be crucial for elucidating whether they act as beneficial or pathological responses in disease and for identifying novel therapeutic targets.

As with many emerging mechanisms in lysosomal damage responses, the precise roles and regulatory processes of PB formation remain to be fully elucidated. For example, comprehensive investigation is needed to evaluate the overall contribution of PBs to cellular homeostasis during lysosomal recovery. While PBs are primarily recognized for their roles in translational control, further research is needed to better understand how they regulate specific RNA populations, influence translational output in support of lysosomal recovery and how RNA metabolism contributes to PB formation. Given their critical role in promoting lysosomal recovery and cell survival, and their close interplay with SGs, PBs likely serve multiple coordinated functions in restoring cellular homeostasis during stress recovery. Moreover, the specific signaling mechanisms that distinguish the recovery phase from acute stress response and that govern PB formation also remain to be defined. Together, these questions highlight the need for future studies to elucidate the molecular mechanisms underlying PB function, formation, and SG-PB coordination during lysosomal stress.

In conclusion, this study defines the acute and recovery stages of lysosomal damage through translational regulation and identifies PB formation as a key feature of the recovery phase, acting in concert with SGs to restore lysosomal function and promote cell survival during stress resolution. These findings provide important insights into the mechanisms governing lysosomal stress responses and their potential dysregulation in pathological conditions.

## Acknowledgments

This work was supported by an NIGMS R35 grant (R35GM154651) to J. Jia. We thank Dr. Roy Parker (University of Colorado Boulder) for providing the DDX6KO and G3BP1/2&DDX6KO cell lines(Ripin et al., 2024). We thank Dr. Ross Buchan (University of Arizona) for his thoughtful and constructive feedback. We thank the AIM Center (University of New Mexico, funded by the NIH AIM COBRE grant P20GM121176) for instrumental support with imaging and data acquisition.

## Author contributions

J. Jia conceptualized the study, designed the experiments, and analyzed the data; J. Jia and J. Duran performed the majority of the experiments; J. Jia supervised the project. L. Chen contributed to the high content microscope experiments; J. Pu and P. Ivanov provided guidance on experimental design. J. Jia wrote the manuscript with input from J. Pu, R. Buchan and P. Ivanov.

---

**Figure S1.**
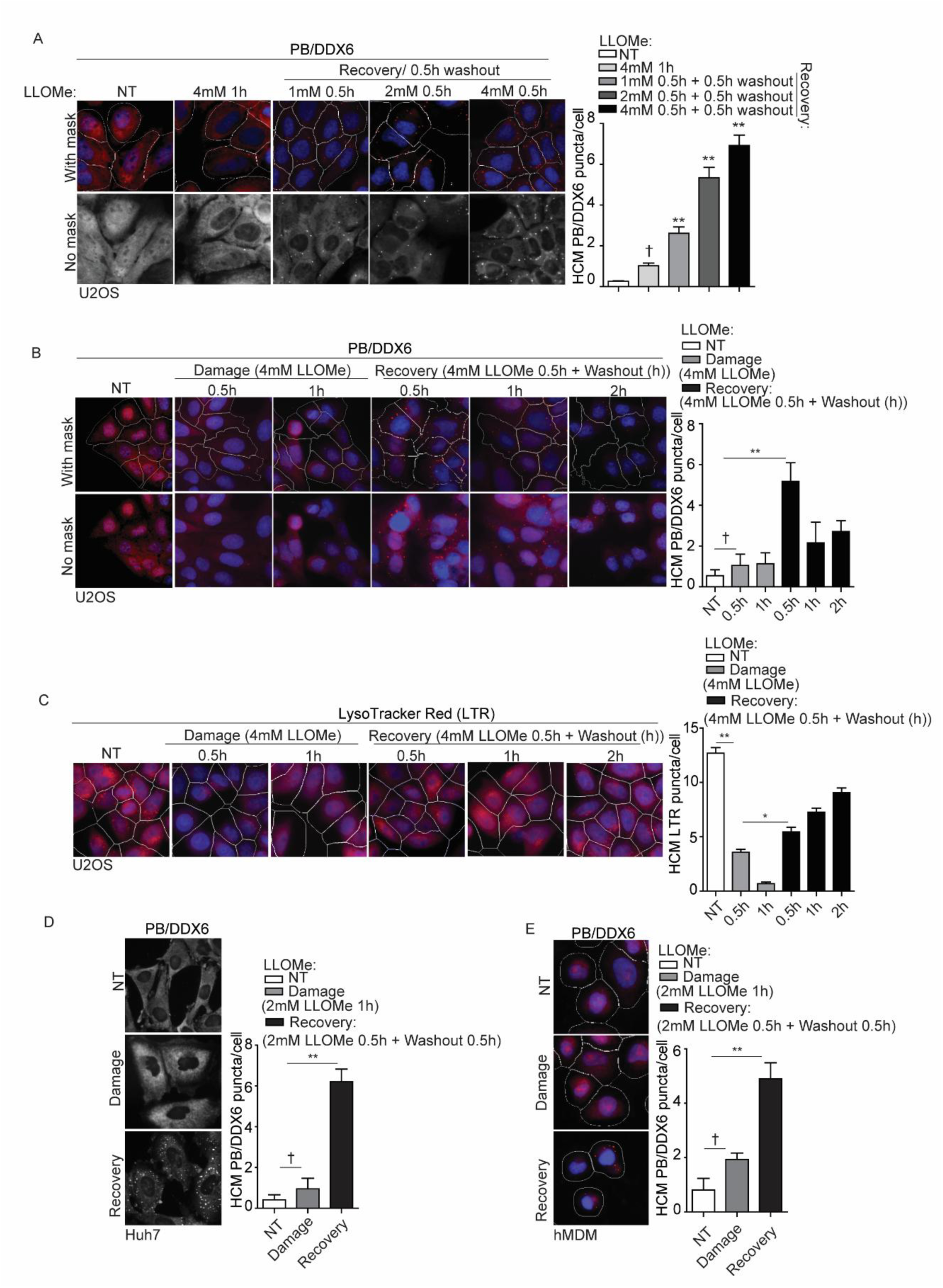
Processing body formation is associated with recovery from lysosomal damage. **(A)** Quantification of the processing body (PB) marker DDX6 by high-content microscopy (HCM) in U2OS cells treated with 4 mM LLOMe for 1 h to induce damage, or with a dose-dependent LLOMe treatment for 30 min, followed by a 30 min recovery after washout. White masks, algorithm-defined cell boundaries; red masks, computer-identified DDX6 puncta. No mask image is shown in grayscale. **(B)** Quantification of the PB marker DDX6 by HCM in U2OS cells treated with 4 mM LLOMe for 30 min or 1 h to induce damage, or with 4 mM LLOMe for 30 min, followed by a time-dependent recovery. White masks, algorithm-defined cell boundaries; red masks, computer-identified DDX6 puncta. **(C)** Status of acidified organelles in U2OS cells assessed by LysoTracker (LTR) staining and HCM during treatment as described in (B). White masks, algorithm-defined cell boundaries; red masks, computer-identified LTR puncta. **(D)** Quantification of the PB marker DDX6 by HCM in Huh7 cells treated with 2 mM LLOMe for 1 h to induce damage, or for 30 min followed by a 30 min recovery after washout. The image is shown in grayscale. **(E)** Quantification of the PB marker DDX6 by HCM in human peripheral blood monocyte-derived macrophages (hMDM) treated with 2 mM LLOMe for 1 h to induce damage, or for 30 min followed by a 30 min recovery after washout. White masks, algorithm-defined cell boundaries; red masks, computer-identified DDX6 puncta. NT, untreated cells. Data, means ± SEM (n = 3); HCM: n ≥ 3 (each experiment: 500 valid cells per well, ≥5 wells/sample). † p ≥ 0.05 (not significant), *p < 0.05, **p < 0.01, ANOVA. See also Figure 1.

**Figure S2.**
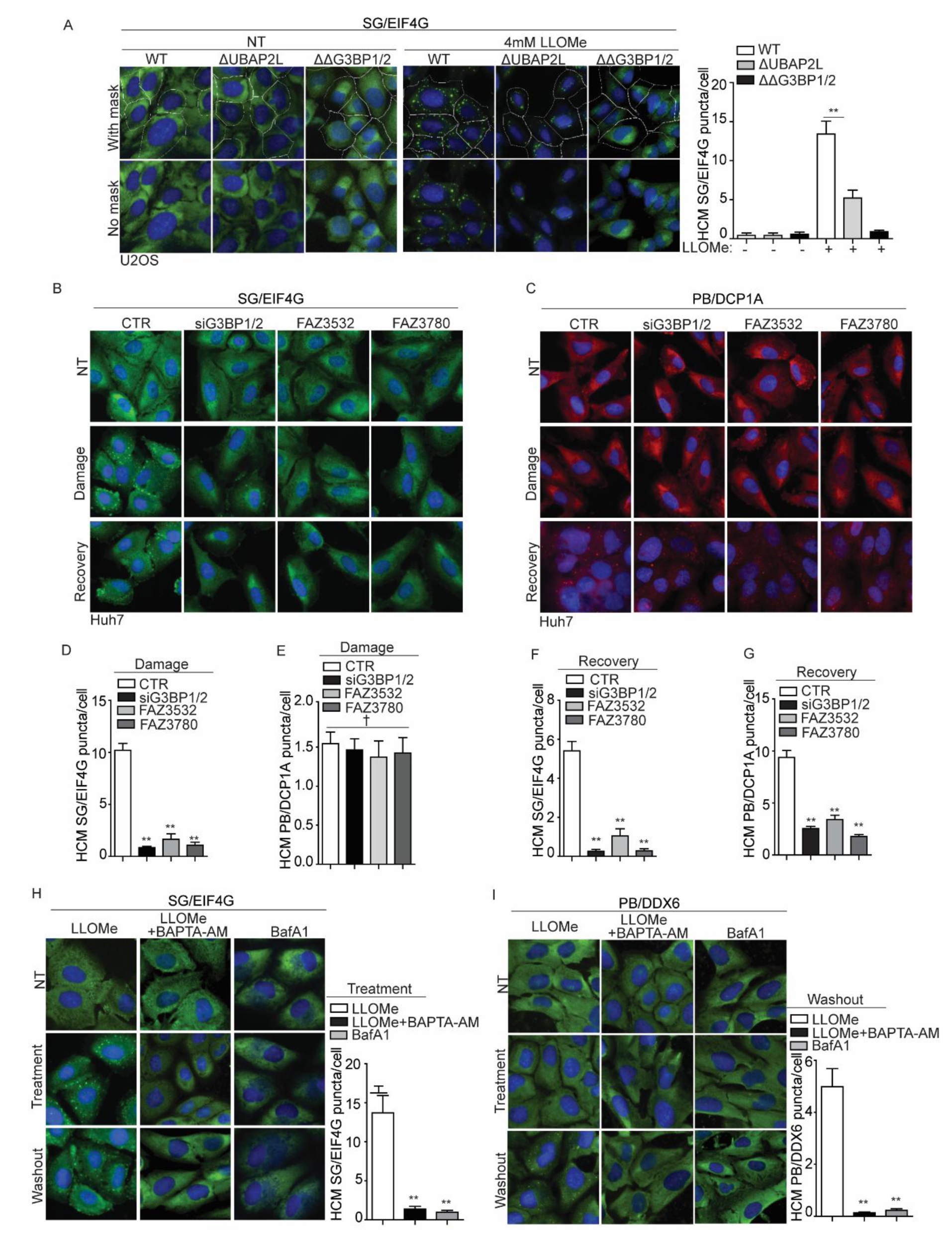
Stress granules facilitate processing body formation during recovery from lysosomal damage. **(A)** Quantification of the SG marker EIF4G by HCM in U2OS wildtype (WT), UBAP2L knockout (ΔUBAP2L), and G3BP1&2 double knockout (ΔΔG3BP1/2) cells treated with 4 mM LLOMe for 1 h. White masks, algorithm-defined cell boundaries; green masks, computer-identified EIF4G puncta. **(B)** Quantification of the stress granule (SG) marker EIF4G by HCM in Huh7 cells transfected with either scrambled siRNA as control (CTR) or siRNA targeting G3BP1 and G3BP2 for knockdown (siG3BP1/2), or treated with SG inhibitors (20 µM FAZ3532/FAZ3780). Cells were treated with 4 mM LLOMe for 1 h to induce damage, or for 30 min followed by a 30 min recovery after washout. **(C)** Quantification of the PB marker DCP1A by HCM in Huh7 cells transfected with either scrambled siRNA as control (CTR) or siRNA targeting G3BP1 and G3BP2 for knockdown (siG3BP1/2) or treated with SG inhibitors. Cells were treated with 4 mM LLOMe for 1 h to induce damage, or for 30 min followed by a 30 min recovery after washout. **(D-G)** Quantification of data shown in (B) and (C). Quantification of the SG marker EIF4G **(H)** and the PB marker DDX6 **(I)** by HCM in U2OS cells. Cells were treated with 4 mM LLOMe with or without 15 µM BAPTA-AM, or with 100 nM Bafilomycin A1(BafA1) for 1 h, followed by a 30 min recovery after washout. NT, untreated cells. Data, means ± SEM (n = 3); HCM: n ≥ 3 (each experiment: 500 valid cells per well, ≥5 wells/sample). † p ≥ 0.05 (not significant), **p < 0.01, ANOVA. See also Figure 2.

**Figure S3.**
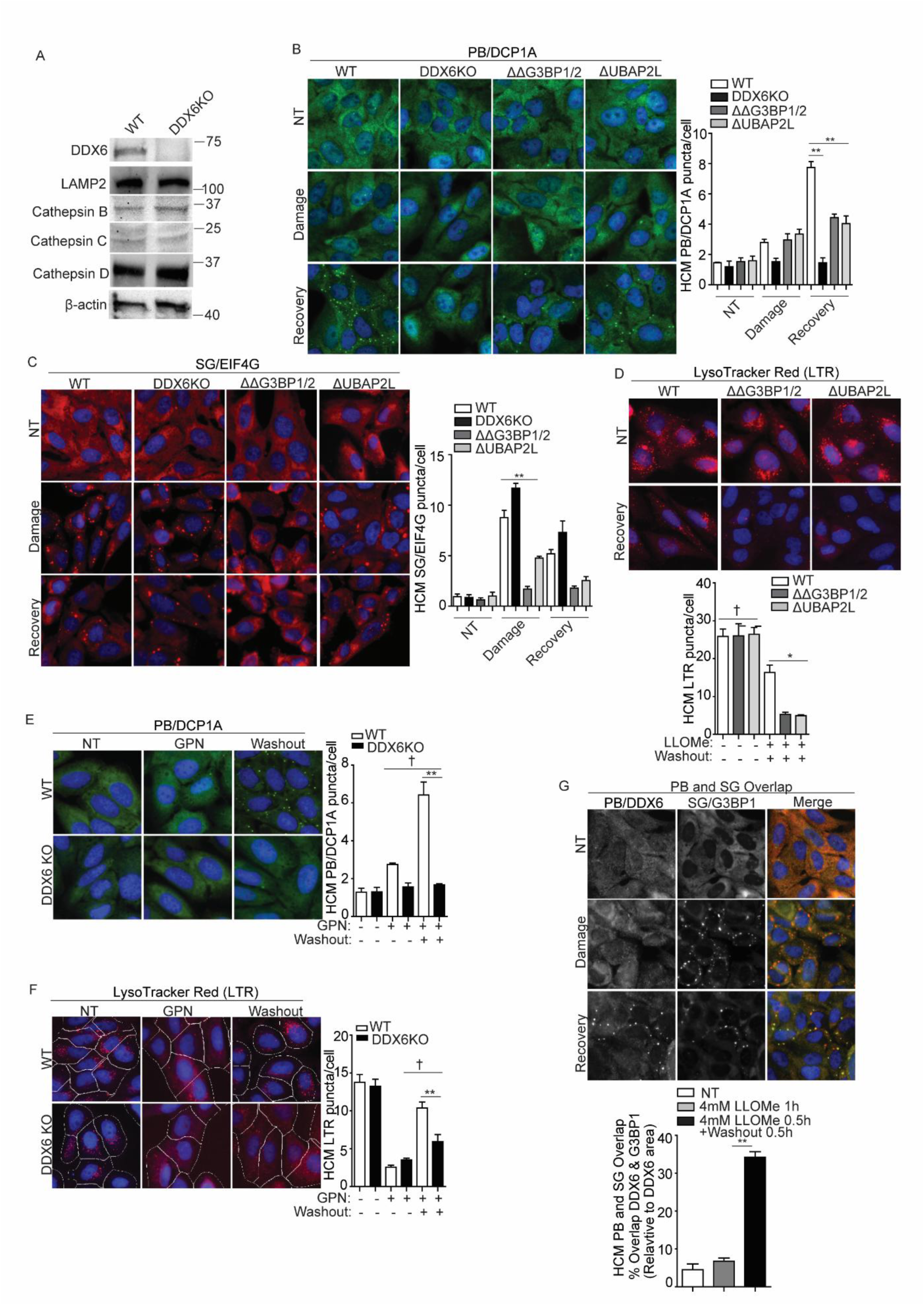
Processing body and stress granule formation during lysosomal recovery is important for maintaining lysosomal quality. **(A)** Immunoblot analysis of the indicated proteins in U2OS wildtype (WT) and DDX6 knockout (DDX6KO) cells. **(B)** Quantification of the PB marker DCP1A (Alexa Fluor 488) by HCM in U2OS wildtype (WT), DDX6 knockout (DDX6KO), G3BP1&2 double knockout (ΔΔG3BP1/2) and UBAP2L knockout (ΔUBAP2L) cells treated with 2 mM LLOMe for 1 h to induce damage or treated for 30 min followed by a 30 min recovery after washout. **(C)** Quantification of the SG marker EIF4G (Alexa Fluor 568) by HCM in U2OS wildtype (WT), DDX6 knockout (DDX6KO), G3BP1&2 double knockout (ΔΔG3BP1/2) and UBAP2L knockout (ΔUBAP2L) cells treated with 4 mM LLOMe for 1 h to induce damage or treated for 30 min followed by a 30 min recovery after washout. **(D)** Status of acidified organelles in U2OS wildtype (WT), G3BP1&2 double knockout (ΔΔG3BP1/2) and UBAP2L knockout (ΔUBAP2L) cells assessed by LysoTracker (LTR) staining and HCM during recovery (2 mM LLOMe for 30 min followed by a 30 min recovery after washout). **(E)** Quantification of the PB marker DCP1A by HCM in U2OS wildtype (WT) and DDX6 knockout (DDX6KO) cells treated with 200 µM GPN for 1 h to induce damage, or for 30 min followed by a 30 min recovery after washout. **(F)** Status of acidified organelles in U2OS wildtype (WT) and DDX6 knockout (DDX6KO) cells assessed by LysoTracker (LTR) staining and HCM during lysosomal damage (200 µM GPN for 1 h) or recovery (200 µM GPN for 30 min followed by a 30 min recovery after washout). White masks, algorithm-defined cell boundaries; red masks, computer-identified LTR puncta. **(G)** HCM quantification of overlaps between the PB marker DDX6 (Alexa Fluor 488) and the SG marker G3BP1 (Alexa Fluor 568) in U2OS cells treated with 4 mM LLOMe for 1 h to induce damage, or for 30 min followed by a 30 min recovery after washout. NT, untreated cells. Data, means ± SEM (n = 3); HCM: n ≥ 3 (each experiment: 500 valid cells per well, ≥5 wells/sample). † p ≥ 0.05 (not significant), *p < 0.05, **p < 0.01, ANOVA. See also Figure 3.

**Figure S4.**
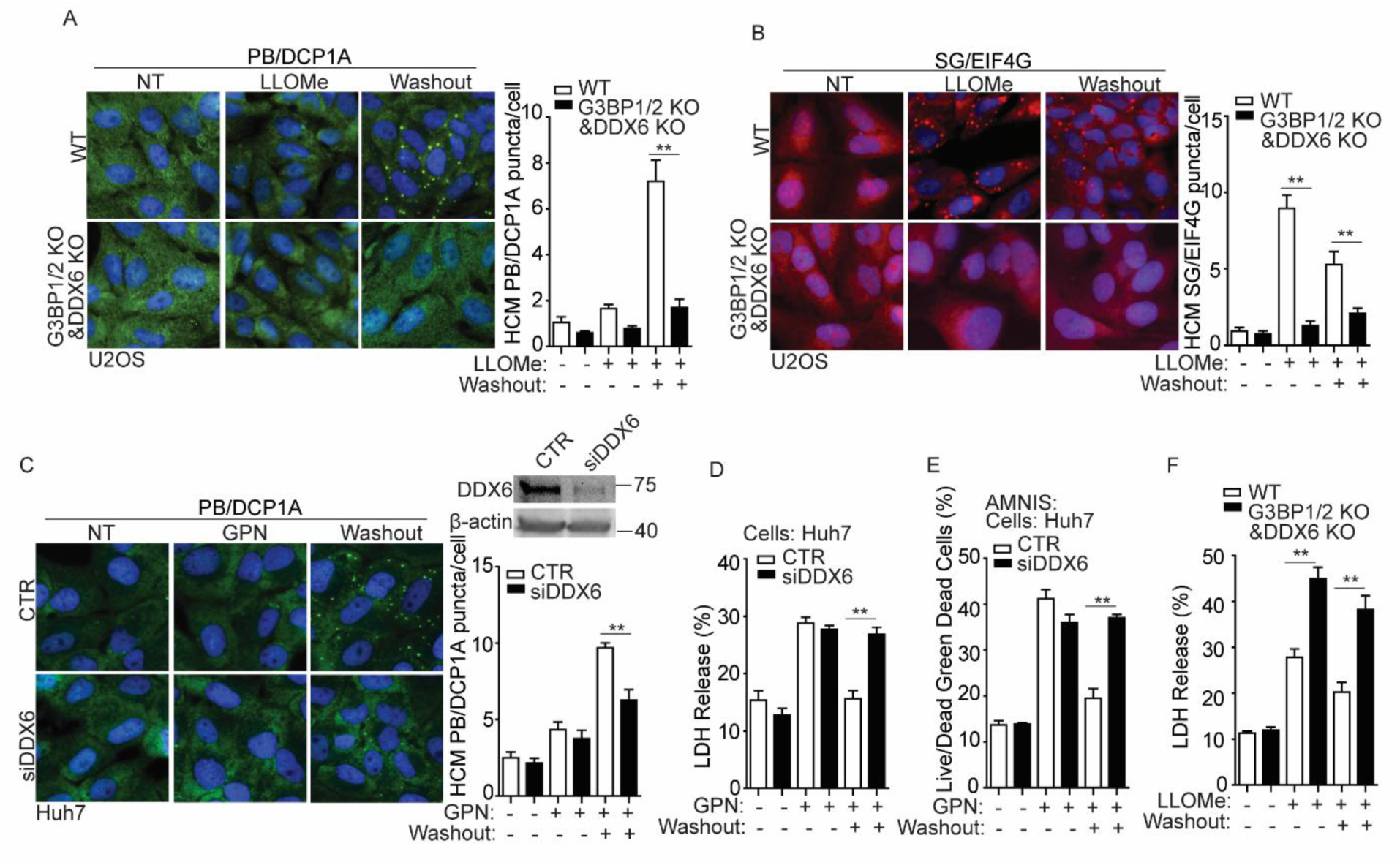
Processing body formation during lysosomal recovery is important for cell survival. (**A)** Quantification of the PB marker DCP1A (Alexa Fluor 488) by HCM in U2OS wildtype (WT) and G3BP1/2 and DDX6 double knockout (G3BP1/2KO&DDX6KO) cells. Cells were treated with 4 mM LLOMe for 1 h to induce damage, or for 30 min followed by a 30 min recovery after washout. (**B)** Quantification of the SG marker EIF4G (Alexa Fluor 568) by HCM in U2OS wildtype (WT) and G3BP1/2 and DDX6 double knockout (G3BP1/2KO&DDX6KO) cells. Cells were treated with 4 mM LLOMe for 1 h to induce damage, or for 30 min followed by a 30 min recovery after washout. **(C)** Quantification of the PB marker DCP1A (Alexa Fluor 488) by HCM in Huh7 cells transfected with either scrambled siRNA as control (CTR) or siRNA targeting DDX6 for knockdown (siDDX6). Cells were treated with 200 µM GPN for 1 h to induce damage, or for 30 min followed by a 30 min recovery after washout. (**D**) Cell death analysis of supernatants from Huh7 cells transfected with either scrambled siRNA as control (CTR) or siRNA targeting DDX6 for knockdown (siDDX6) by an LDH release assay. Cells were treated as described in (C). (**E**) Quantification of cell death using the AMNIS system and Live/Dead^TM^ stain kit in Huh7 cells transfected with either scrambled siRNA as control (CTR) or siRNA targeting DDX6 for knockdown (siDDX6). Cells were treated as described in (C) and then stained using Live/Dead^TM^ stain kit (ThermoFisher). (**F**) Cell death analysis of supernatants from U2OS wildtype (WT) and G3BP1/2 and DDX6 double knockout (G3BP1/2KO&DDX6KO) cells by a LDH release assay. Cells were treated as described in (A). NT, untreated cells. Data, means ± SEM (n = 3); HCM: n ≥ 3 (each experiment: 500 valid cells per well, ≥5 wells/sample). **p < 0.01, ANOVA. See also Figure 4.

## MATERIALS AND METHODS

### Antibodies and reagents

Antibodies from Cell Signaling Technology were Phospho-eIF2α (Ser51)(1:1000 for WB), eIF2α (1:1000 for WB), G3BP2 (1:1000 for WB), Cathepsin B (1:1000 for WB), Cathepsin D (1:1000 for WB). Antibodies from Proteintech: EIF4G1 (15704-1-AP) (1:200 for IF), Cathepsin C (1:1000 for WB), LSM14A (1:200 for IF). Antibodies from Abnova: DCP1A (1:200 for IF). Antibodies from MBL: DDX6 (1:200 for IF) (1:1000 for WB). Antibodies from BioLegend: Galectin-3 (1:1000 for WB; 1:500 for IF). G3BP1(PA5-29455, 1:1000 for WB, 1:200 for IF), Alexa Fluor 488, 568, 647 (1:500 for IF) and secondary antibodies from ThermoFisher Scientific. Other antibodies used in this study were from the following sources: beta-Actin (C4)(1:1000 for WB) from Santa Cruz Biotechnology; LAMP2 (H4B4)(1:500 for IF) from DSHB of University of Iowa; UBAP2L (1:1000 for WB) from Bethyl.

Leu-Leu-methyl ester hydrobromide (LLOMe) from Sigma Aldrich. Gly-Phe-β-naphthylamide (GPN) from Cayman. Reagents from ThermoFisher were Hoechst 33342, LIVE/DEAD™ Fixable Green Dead Cell Stain Kit, Click-iT™ Plus OPP Alexa Fluor™ 488 Protein Synthesis Assay Kit, Lipofectamine RNAiMAX Transfection Reagent, BP/LR Clonase Plus Enzyme Mix, Prolong Gold Antifade Mountant with DAPI, Human M-CSF Recombinant Protein, DMEM, Opti-MEM Reduced Serum Media, EBSS, PBS, Penicillin-Streptomycin, Fetal Bovine Serum, NP40 Cell Lysis Buffer and LysoTracker Red DND-99. Reagents from Promega were CytoTox 96® Non-Radioactive Cytotoxicity Assay, FuGENE HD Transfection Reagent and ProFection Mammalian Transfection System. Other reagents used in this study were from the following sources: Protease Inhibitor from Roche (11697498001); FAZ3532 and FAZ3780 from MedChemExpress (HY-162288 and HY-162289); BAPTA-AM from ThermoFisher (B1205); Anti-HA Magnetic Beads from ThermoFisher (88836); Bafilomycin A1 from InvivoGen.

### Cells and cell lines

U2OS cells were from American Type Culture Collection. Huh7 cells are from Rocky Mountain laboratories. Human peripheral blood monocyte-derived macrophages (hMDM) were derived from peripheral blood mononuclear cells (PBMCs) isolated from venipuncture blood from anonymous donors, details below. Knockdown cell lines were generated by small interfering RNAs (siRNAs) from GE Dharmacon (siGENOME SMART pool), details below. U2OS wildtype (WT), G3BP1&2 double knockout (ΔΔG3BP1/2) and UBAP2L knockout (ΔUBAP2L) cells were from Dr. Pavel Ivanov (Brigham and Women’s Hospital and Harvard Medical School, Boston, MA). DDX6 knockout (DDX6KO) and G3BP1/2 and DDX6 double knockout (G3BP1/2KO&DDX6KO) cells were from Dr. Roy Parker (University of Colorado Boulder)

### Cultured human peripheral blood monocyte-derived macrophages

40-50 mL of venipuncture blood was collected from healthy, consenting adult volunteers at the Vitalant Blood Donation Center (Albuquerque, NM). Blood from individual donors (10 mL vacutainer tubes) was placed into two 50 mL conical tubes and the volume was brought to 50 mL with sterile 1 X PBS followed by mixing inversely. 25 mL of the blood mix were carefully layered onto 20 mL of Ficoll (Sigma, #1077) in separate conical tubes and centrifuged at 2000 rpm for 30 min at 22 °C. The buffer layer containing human peripheral blood monocytes (PBMCs) was removed, pooled, washed with 1X PBS twice, and resuspended in 20 mL RPMI media with 10 % human AB serum and Primocin. PBMCs were cultured in RPMI 1640 with GlutaMAX and HEPES (Gibco), 20 % FBS, and 200 ng/mL Human M-CSF Recombinant Protein (ThermoFisher). Six days after the initial isolation, differentiated macrophages were detached in 0.25 % Trypsin-EDTA (Gibco) and seeded for experiments.

### Plasmids, siRNAs, and transfection

Plasmids used in this study, e.g., G3BP1 and DDX6 cloned into pDONR221 using BP cloning, and expression vectors were made utilizing LR cloning (Gateway, ThermoFisher) in appropriate pDEST vectors for immunoprecipitation assay. Small interfering RNAs (siRNAs) were from Horizon Discovery (siGENOME SMART pool), DDX6 (L-006371-00-0005), G3BP1(L-012099-00-0005), G3BP2 (L-015329-01-0005). Plasmid transfections were performed using the ProFection Mammalian Transfection System, FuGENE® HD Transfection Reagent (Promega), or Lipofectamine 2000 Transfection Reagent (ThermoFisher). siRNAs were delivered into cells using Lipofectamine RNAiMAX (ThermoFisher).

### High content microscopy (HCM) analysis

Cells in 96 well plates were fixed in 4 % paraformaldehyde for 5 min. Cells were then permeabilized with 0.1 % saponin in 3 % Bovine serum albumin (BSA) for 30 min followed by incubation with primary antibodies for 2 h and secondary antibodies for 1 h. Hoechst 33342 staining was performed for 3 min. HCM with automated image acquisition and quantification was carried out using a Cellomics HCS scanner and iDEV software (ThermoFisher). Automated epifluorescence image collection was performed for a minimum of 500 cells per well. Epifluorescence images were machine analyzed using preset scanning parameters and object mask definitions. Hoechst 33342 staining was used for autofocus and to automatically define cellular outlines based on background staining of the cytoplasm. Primary objects were cells, and regions of interest (ROI) or targets were algorithm-defined by shape/segmentation, maximum/minimum average intensity, total area and total intensity, to automatically identify puncta or other profiles within valid primary objects. All data collection, processing (object, ROI, and target mask assignments) and analyses were computer driven independently of human operators. HCM provides variable statistics since it does not rely on parametric reporting cells as positive or negative for a certain marker above or below a puncta number threshold.

### Protein translation assay

Cells in 96 well plates were subjected to indicated treatment and then stained with O-propargyl-puromycin (OPP) using the Click-iT Plus OPP Alexa Fluor 488 protein synthesis assay kit (Thermo Fisher Scientific) in accordance with the manufacturer’s guidelines. Cells were scanned by HCM, described below. We quantified by HCM the overall fluorescence intensity in cells of fluorescently labeled OPP. OPP is an alkyne analog of puromycin that following incorporation into polypeptides can be fluorescently labeled by Alexa 488 picolyl azide through a chemoselective ligation or “click” reaction, occurring between the picolyl azide dye and the OPP alkyne.

### LDH release assay

Each well of a 96-well plate was initially plated with 20,000 cells. Cells were treated with lysosomal damaging agents as indicated. Following this, the supernatant was measured for LDH (Lactate dehydrogenase) release using the kit of CytoTox 96® Non-Radioactive Cytotoxicity Assay (Promega, G1780), according to the manufacturer’s instructions.

### Amnis flow cytometry analysis

Cells after treatment were washed with 3 % BSA in PBS supplemented with 0.1 % of NaN3 before staining. Cells were stained using LIVE/DEAD™ Fixable Green Dead Cell Stain Kit (ThermoFisher) following the manufacturer’s instructions. After staining, cells were then resuspended with 3 % BSA in PBS supplemented with 0.1 % of NaN3 until acquisition on Amins ImageStreamx MKII (ISx, EMD Millipore, Seattle, WA, USA).

### LysoTracker assay

LysoTracker (LTR) Staining Solution was prepared by freshly diluting 2 μL of LTR stock solution (1 mM LysoTracker Red DND-99; Sigma Aldrich, L7528) in 1 mL of medium. 10 μL of Lyso-Tracker Staining Solution was added to 90 μL of medium each well in 96 well plates (final volume 100 μL per well, final concentration 0.2 μM LTR) and adherent cells incubated at 37°C for 30 min protected from light. Wells were rinsed gently by 1 × PBS and fixed in 4% paraformaldehyde for 2 min. Wells were washed once in 1 × PBS and nuclei stained with Hoechst 33342 for 2 min before analyzing the plates by HCM.

### Extraction of cytosol

Cytosol was extracted using a protocol adapted from previous studies(Appelqvist et al., 2011; Eriksson et al., 2020), using the cholesterol-solubilizing agent digitonin (Sigma-Aldrich). At low concentrations, digitonin selectively permeabilizes the cholesterol-rich plasma membrane while preserving the integrity of intracellular membranes with lower cholesterol content. Cell culture dishes were incubated on ice with gentle rocking for 15 min in extraction buffer (250 mM sucrose, 20 mM HEPES, 10 mM KCL, 1.5 mM MgCL₂, 1 mM EGTA, 1 mM EDTA, pH 7.5) containing 25 μg/mL digitonin. Following incubation, the extraction buffer—containing the cytosolic fraction— was collected and kept on ice. For western blot analysis, cytosolic proteins were precipitated by adding an equal volume of methanol and one-third volume of chloroform, followed by centrifugation. The resulting pellets were resuspended in Laemmli buffer.

### Immunofluorescence confocal microscopy analysis

Cells were plated onto coverslips in 6-well plates. After treatment, cells were fixed in 4 % paraformaldehyde for 5 min followed by permeabilization with 0.1 % saponin in 3 % BSA for 30 min. Cells were then incubated with primary antibodies for 2 h and appropriate secondary antibodies Alexa Fluor 488 or 568 or 647 (ThermoFisher) for 1 h at room temperature. Coverslips were mounted using Prolong Gold Antifade Mountant (ThermoFisher). Images were acquired using a confocal microscope (META; Carl Zeiss) equipped with a 63 3/1.4 NA oil objective, camera (LSM META; Carl Zeiss), and AIM software (Carl Zeiss).

### Quantification and statistical analysis

Data in this study are presented as means ± SEM (n ≥ 3). Data were analyzed with either analysis of variance (ANOVA) with Tukey’s HSD post-hoc test, or a two-tailed Student’s t test. For HCM, n ≥ 3 includes in each independent experiment: 500 valid primary objects/cells per well, from ≥ 5 wells per plate per sample. Results are presented as mean ± SEM. Statistical significance was determined using ANOVA, with p < 0.05 considered statistically significant. Statistical significance was defined as: † (not significant) p ≥ 0.05 and *p < 0.05, **p < 0.01.

